# Exploring the molecular function of metabolites identified in Elite Controllers and their role in epithelial integrity and immune regulation

**DOI:** 10.64898/2026.07.29.741523

**Authors:** Itziar Chapartegui Gonzalez, Aswathy Narayanan, Rafael Ceña Diez, Anders Sönnerborg, Shilpa Ray

## Abstract

Despite advances in treatment, HIV-1 infection continues to remain a major global health challenge, prompting ongoing efforts to understand the mechanisms that enable natural viral suppression and immune control. Elite controllers (ECs), a rare subset of PLWH individuals, naturally suppress HIV-1 replication without antiretroviral therapy, highlighting the importance of host-related factors in viral control. Understanding the mechanisms underlying this unique phenotype is crucial for developing novel therapeutic strategies. Previous studies from our group identified certain EC-specific metabolites, called dipeptides (DPs), and investigated their antiviral properties. We hypothesize that these dipeptides may potentially affect epithelial barrier integrity by modulating the expression of tight junction proteins, which in turn influences the mucosal barrier function, a key factor in HIV-1 pathogenesis. Therefore, in this study we investigated the impact of ten EC-specific DPs on tight junction (TJ) gene and protein expression in epithelial models derived from the female reproductive and gastrointestinal tracts, where we observed enhanced expression of different TJ genes (*CLDN1, CLDN3, CLDN4, CLDN7, CLDN14, TJP1, TJP2, OCLN*) and proteins (CLDN1, CLDN7, and CLDN14), suggesting the potential influence of these dipeptides on epithelial barrier function. Furthermore, we also examined different proteomic profiles between dipeptide (WG)-treated HeLa CD4^+^ CCR5^+^ cells compared with the untreated ones, and observed significantly reduced abundance of pro-inflammatory proteins, such as RELB Proto-Oncogene (RELB), TNF-α-induced protein 1 (TNFAIP1), TNF receptor superfamily member 1A (TNFRSF1A), and IL-32, in dipeptide-treated cells; and increased expression of proteins associated with tissue homeostasis (SMAD family member 5 [SMAD5]), cellular proliferation (transforming growth factor β receptor 3 [TGFBR3]), and epithelial integrity, like CD81. Interestingly, KEGG analysis revealed possible attenuation of NF-κB, MAPK, TNF, and JAK-STAT signaling pathways, along with the enrichment of mTOR and PI3K-AKT pathways in treated HeLa CD4^+^ CCR5^+^ cells. Overall, this study investigated the potential interplay between tight junction proteins and key signaling pathways involved in maintaining epithelial barrier integrity and modulating immune activation, potentially contributing to both HIV-1 control and to the chronic inflammation associated with infection.

## INTRODUCTION

Human Immunodeficiency Virus type 1 (HIV-1), a pathogenic virus that infects the body’s immune system specifically targeting CD4^+^ T-helper lymphocytes, remains a significant public health issue globally, with approximately 39.9 million people living with HIV-1 (PLWH) and over 630,000 fatalities reported in 2023, despite advances in antiretroviral therapies (ART) (The World Health Organization, 2024). Although ART can effectively suppress viral replication, HIV-1 can persist in several cellular and tissue reservoirs throughout the body, where the virus can remain latent for years, contributing to disease progression and persistence, accompanied by intensive immune activation (Walker and Yu, 2013; Woldemeskel et al., 2020). Notably, this immune activation is strongly related to microbial translocation from the gut to the blood through damaged gut-blood barrier (Troseid et al., 2011), since modern combination ART fails to fully restore the mucosal barrier defects or eliminate chronic inflammation (Cassol et al., 2010; Merlini et al., 2011; Nowak et al., 2012; Vesterbacka et al., 2013), leaving patients at higher risk of non-communicable comorbidities (Troseid et al., 2011; Nowak et al., 2012; Antony et al., 2020).

Interestingly, among the PLWH, a rare subset of individuals (<1%), known as elite controllers (ECs), can maintain undetectable viral loads in blood plasma and control the HIV-1 replication, while also keeping stable CD4^+^ T-cell counts even without ART (Walker and Yu, 2013; Gebara et al., 2019; Bai et al., 2024). While previous studies evaluated whether infection with attenuated or replication-defective viral strains was responsible for this phenotype, current studies suggest that the EC phenotype results primarily from host-related factors, including potent immune responses and favorable genetic determinants (Walker and Yu, 2013; Salgado et al., 2024). ECs are characterized by highly effective HIV-specific CD8^+^ T-cell responses, reduced T-cell exhaustion, and genetic factors that promote viral control (Walker and Yu, 2013; Bai et al., 2024). While CD8^+^ T-cell responses are the dominant feature of immune control in EC, other host genetic factors, including specific protective HLA class I alleles, particularly HLA-B57 and HLA-B27 (Walker and Yu, 2013), or polymorphisms in genes such as the CCR5 (CC-chemokine receptor 5), also contribute to the durable control of HIV-1 infection. However, these genetic factors alone are neither necessary nor sufficient for EC phenotype, highlighting its multifactorial nature (Walker and Yu, 2013; Leon et al., 2016; Claireaux et al., 2022).

Beyond the distinct microbial composition observed in ECs (Vesterbacka et al., 2017), maintenance of epithelial barrier integrity is another key factor influencing HIV-1 pathogenesis. Tight junctions (TJs) regulate epithelial permeability and prevent microbial translocation, while microbiota-derived metabolites, such as short-chain fatty acids (SCFAs), support barrier function and mucosal homeostasis (Ganieva et al., 2022; Moonwiriyakit et al., 2023). During HIV-1 infection, disruption of the gut barrier leads to increased permeability due to compromised tight junction integrity, resulting in barrier dysfunction. This promotes microbial translocation (“leaky gut”), allowing not only bacteria but also their products (i.e., LPS) into systemic circulation, which subsequently causes chronic immune activation and increased risk of opportunistic infections and other comorbidities (Klatt et al., 2013). This process contributes to impair CD4⁺ T-cell recovery during ART and persistence of viral reservoirs (Tincati et al., 2016). Although most studies have focused on the gastrointestinal tract, barrier integrity is equally important in the female reproductive tract. *Lactobacillus*-dominated vaginal microbiota strengthens mucosal barrier function and is associated with reduced HIV-1 acquisition risk (Hearps et al., 2017; Delgado-Diaz et al., 2022). Furthermore, crosstalk between the gut and vaginal microbiota suggests that similar barrier-protective mechanisms may influence local immunity and HIV-1 transmission in women (Amabebe and Anumba, 2020).

Recent metabolomic analyses from our group have identified an enrichment of antiviral dipeptides (DPs) in ECs when compared with viremic progressors and non-infected individuals, suggesting that these microbiota-associated metabolites may contribute to immune regulation and HIV-1 control (Sperk et al., 2021). However, whether these antiviral DPs (WG, VQ) directly influence epithelial barrier integrity and TJ regulation remains unknown, highlighting the need to investigate their potential role in maintaining mucosal barrier function. To address this, we explored the effects of ten EC-specific DPs (Sperk et al., 2021) on tight-junction expression in epithelial cell models derived from the gastrointestinal and female reproductive tracts. Through gene-expression and protein-level analyses, we aimed to determine whether these EC-specific metabolites, which were subsequently shown to possess antiviral ability, can be crucial for EC-status of individuals (Ceña-Diez et al., 2022, 2023) by contributing to epithelial barrier regulation. Given this premise, we also conducted a proteomic analysis where HeLa CD4^+^ CCR5^+^ cells were treated with different doses of WG, one of the described EC-specific metabolites (Ceña-Diez et al., 2023). In this study, we therefore compared the proteins expressed between dipeptide (WG)-treated cells with the untreated ones and observed differential expression of several proteins associated with inflammatory responses, immune signaling, and cellular stress pathways, hinting at a potential immunomodulatory role of the metabolite. This approach provides an initial mechanistic insight into how EC-specific metabolites might influence host-cell responses beyond their previously described antiviral activity. Overall, this study hints at the potential role of EC-specific fecal metabolites as barrier-protective, signaling-modulatory agents relevant to HIV-1 persistence and transmission, thus potentially leading to the future use of these DPs as clinical prophylactic agents against HIV-1 in the broader PLWH population.

## MATERIALS AND METHODS

### Cell culture

HEC-1A cells (HTB-112™), derived from human endometrial adenocarcinoma; VK2/E6E7 (CRL-2616™), human vaginal epithelial cell line; and HCT-116 (CCL-247™) derived from human colorectal carcinoma were purchased from ATCC.

HEC-1A cells were maintained in McCoy’s 5A modified medium (GIBCO 26600-023, ThermoScientific); VK2 in Keratinocyte-Serum Free medium (GIBCO-BRL 17005-042, ThermoScientific) supplemented with 0.1 ng/ml human recombinant EGF, 0.05 mg/ml bovine pituitary extract, and additional calcium chloride final concentration 0.4 mM; and HCT-116 in DMEM GlutaMax (GIBCO 31966-047, ThermoScientific). All three media were supplemented with antibiotics (100 µg/mL streptomycin, 100 U/mL penicillin [Sigma]), while 10% FBS (GIBCO A5670801, ThermoScientific) was only added to McCoy’s and DMEM media.

### Cell treatments

For gene expression, cells were seeded at 1×10^5^ cells/well confluency in 48-well cells (Nunc™, ThermoScientific). The cells were treated with the different dipeptides (Table 1) (Sperk et al., 2021) at a final concentration of 5 mM for 2 h, 6 h, 24 h, or 48 h incubation periods. Following the incubation times, the cells were stored in DNA/RNA Shield (R1100; Zymo Research) until extraction. Additionally, lactic acid (LA, at a non-cytotoxic percentage, such as 0,075% and 0,15%), LPS (0.5 ng/mL and 1 ng/mL), and vitamin D (10 nM and 100 nM) were also added to cells for experimental controls. LA treatment was performed with slight protocol modification, where the cells were treated for 1 h with subsequent fresh media replacement for the rest of incubation period to avoid cytotoxicity. For all the treatments and timepoints, untreated cells were considered as control.

**Table 1:** List of dipeptides used in this study.

| Code | Dipeptides |
| --- | --- |
| P1 | WG-OH |
| P2 | VQ-OH |
| P3 | WG-NH<sub>2</sub> |
| P4 | VQ-NH<sub>2</sub> |
| P5 | LQ-NH<sub>2</sub> |
| P6 | AL-NH<sub>2</sub> |
| P7 | TF-NH<sub>2</sub> |
| P8 | LA-NH<sub>2</sub> |
| P9 | GV-NH<sub>2</sub> |
| P10 | WG-SO<sub>4</sub> |

### Cell Cytotoxicity

All the cell treatments described above were tested for cytotoxicity before analysis. Cells were seeded in 96-well black plates at a confluency of 1×104 cells/well. Following the treatments and incubations, an equivalent volume of CellTiter-Glo® 2.0 Cell Viability Assay (G9242, Promega) was added to each well and incubated for 10 min at room temperature (RT). The luminescence value of each well was read on a Spark Microplate Reader (Tecan). The % of cell survival for each sample was calculated as a % of the luminescence value considering the non-treated cells (control) as 100%.

### RNA extraction and cDNA synthesis

RNA was extracted using Quick-RNA Miniprep Plus Kit (R1058, Zymo Research), according to the manufacturer’s instructions. RNA concentration and purity were measured using a NanoDrop ND-1000 Spectrophotometer (ThermoScientific) and stored at −80°C for future use.

The different RNA samples were used for cDNA synthesis using High-Capacity cDNA Reverse Transcription kit (ThermoScientific), following the manufacturer’s instructions, with the temperature conditions: 25°C for 10 min, 37°C for 120 min, 85°C for 5 min, 4°C hold. The synthesized cDNA was stored at −20°C for further use.

### Expression quantification by RT-PCR analysis

Quantitative PCR (qPCR) was used to analyze the expression of the tight-junction protein genes (*CLDN1, CLDN3, CLDN4, CLDN7, CLDN14, OCLN, TJP1, TJP2*) for the different cell lines after the various treatments, as described earlier. Gene expression quantification was performed using KAPA SYBR® FAST qPCR Master Mix (KK4602, Roche) following the manufacturer’s instructions in a Bio-Rad CFX96 thermocycler, considering the untreated samples as the control for expression in each condition. Expression was normalized with the housekeeping gene *β-actin* for each condition, following the ΔΔ*Ct* method (Livak and Schmittgen, 2001). All primers used for expression are mentioned in Table 2. The PCR cycling program was set as follows: initial heat activation step 50°C 2 min and 92°C 10 min; two-step 40 cycles of 15 s 95°C and 60 s 60°C. Additionally, to compare the baseline expression of the genes among the cell lines or at different timepoints, the relative expression was considered as follows: 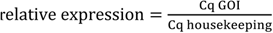 (GOI: Gen Of Interest), in which higher values represent lower expression. Ordinary two-way ANOVA test was conducted to establish significant differences among different conditions, and significance was considered *p* < 0.05.

**Table 2:**
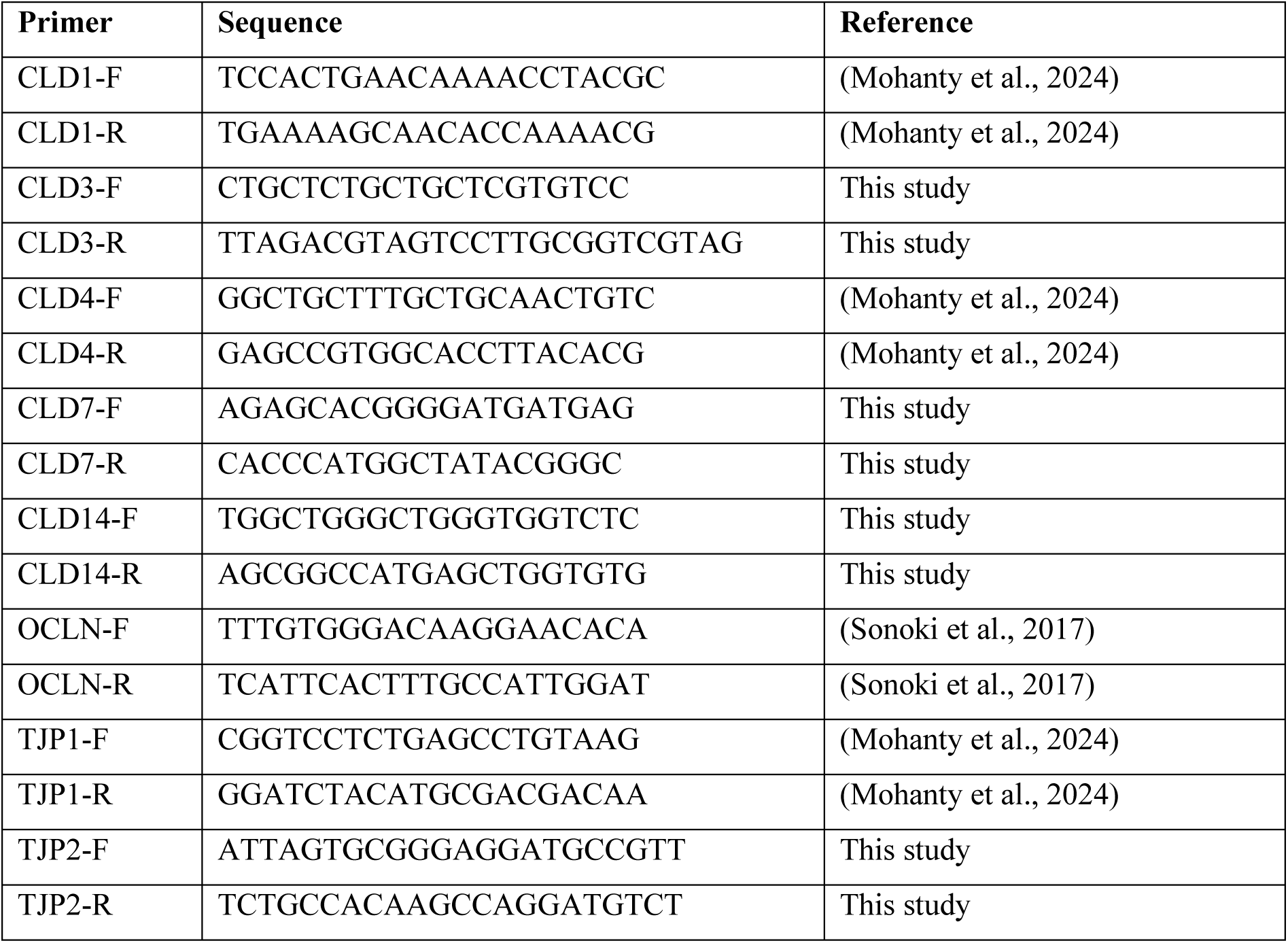
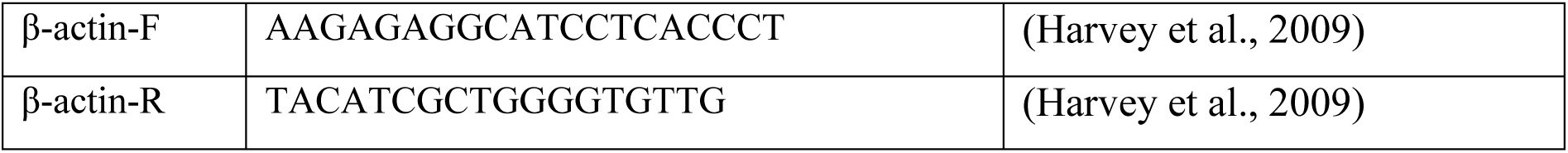
List of primers used in this study.

### Immunofluorescence

For immunofluorescence analysis, cells were seeded in 8-well chambers at a confluency of 1×10^5^ cells/well. The cells were treated, as previously described, in different conditions for either 2-6 h for HCT-116 or 24 h for VK2. Following the incubation, the cells were washed 3 times with PBS and fixed with 4% PFA (28908, ThermoScientific) at RT. Cells were lysed with 0.1% TritonX100 for 10 min at RT and subsequently stained overnight with the primary antibodies (Table 3). Afterwards, cells were incubated with secondary antibodies and phalloidin for actin staining (Alexa Fluor 488 phalloidin [A12379, Invitrogen], Alexa Fluor 594 phalloidin [A12381, Invitrogen]) for 1 h each at RT. The slides were mounted using ProLong™ Diamond Antifade Mountant with DAPI (P36966, Invitrogen), followed by visualization in Nikon A1R confocal microscope, and analyzed using the Image J software for MFI quantification (version 1.52h). The region of interest (ROI) was selected, and the MFI was quantified in all the channels. The *raw* MFI of each claudin was obtained and it was also normalized based on the DAPI MFI 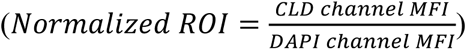, to account for the number of cells in each ROI.

**Table 3:** List of antibodies used in this study.

| Antibody | Company | Reference |
| --- | --- | --- |
| anti-CLDN1 | Invitrogen | 37-4900 |
| anti-CLDN7 | Invitrogen | 37-4800 |
| anti-CLDN14 | Invitrogen | 36-4200 |
| anti-mouse IgG | Invitrogen | A32728 |
| anti-rabbit IgG | Invitrogen | A32731 |

### Proteomics data processing and analysis

Protein identification and quantification were performed as previously described (Ceña-Diez et al., 2023). The raw protein data was normalized using R package NormalyzerDE v1.20.0 (Willforss et al., 2019). Quantile normalized data was used for further analysis. Differential abundance analysis was performed using R package Linear models for microarray (Limma v3.42.2) (Ritchie et al., 2015) with Benjamini-Hochberg correction for multiple hypotheses to provide adjusted p-value for the comparison between the groups. Pathway analysis of differentially expressed proteins was performed using Database for Annotation, Visualization and Integrated Discovery (DAVID) 6.8 (https://davidbioinformatics.nih.gov/) with default settings. Further, visualization of differentially expressed proteins and pathways was achieved using bubble plots and volcano plots using R package ggplot2 v3.5.1 (Wickham, 2016).

### *In silico* predictions

To predict the protein-protein interaction from the different claudins and tight-junction proteins in human cells, STRING database version 12.0 [http://string-db.org] was used with the default setting of medium confidence required score (0.4) (Szklarczyk et al., 2021). Individual STRING interaction networks were generated for each gene evaluated by qPCR and imported into Cytoscape (version 3.10.4). The individual networks were merged into a consensus network and subsequently filtered to retain the nodes shared across the individual networks with a degree 2, restricting the analysis to the most highly interconnected proteins. In the merged network, node color and size were mapped to degree (number of direct interactions), edge width was mapped to the text-mining score, and edge and label color were mapped to the interaction score (combined confidence score for co-expression, experimental, databases, and text-mining), all calculated in Cytoscape.

### Statistics

All statistical analysis was performed using the GraphPad Prism software (version 10.0.2) and *p* values of <0.05 were considered statistically significant. Quantitative data are presented as mean with standard error (SEM). Significant differences in gene expression were assessed via ordinary two-way ANOVA comparison test. Significant differences in protein expression were assessed via ordinary one-way ANOVA comparison test.

## RESULTS

### Baseline expression of tight junction genes in epithelial cells

In our current study, we analyzed the role of some metabolites in two cell lines derived from the female reproductive tract (endometrial cells, HEC-1A, and vaginal epithelial cells, VK-2) and one from the gastrointestinal tract (colon cancer cell line, HCT-116); that were exposed to 10 distinct dipeptides (listed in Table 1) previously identified in elite controllers (EC) from our published findings (Sperk et al., 2021).

Dipeptide (DP) concentration used was determined after cytotoxicity evaluation (Fig. S1). Interestingly, differential gene expression patterns were observed between the two female reproductive tract cell lines (Fig. S2). To assess baseline gene expression across the cell lines, relative expression levels were quantified, with higher values indicating reduced expression (see Methods). For instance, *occludin* (*OCLN*) exhibited no detectable expression in HEC-1A but was expressed in VK2 (Fig. S2). Furthermore, a comparative analysis revealed distinct expression patterns between the two cell lines (Fig. S2), as our findings indicate that *claudin-1* (*CLDN1*), *claudin-3* (*CLDN3*)*, claudin-4* (*CLDN4*)*, TJP1*, and *TJP2* showed higher expression in HEC-1A compared with VK2 at 24 h incubation, whereas *claudin-7* (*CLDN7*) exhibited higher expression in VK2 compared with HEC-1A at 48 h incubation. Overall, the time-dependent expression of the TJ genes was more consistent and stable in HEC-1A than in VK2, which exhibited a general reduced expression of the genes at 24 h compared with both 6 h and 48 h. In VK2, *OCLN* exhibited the highest expression among all the tested TJ genes at 6 h, in contrast with *claudin-14* (*CLDN14*) that had the lowest at the same time. This highlights the tissue-specific expression of TJ genes.

### Induced tight junction gene expression in epithelial cells due to EC-specific dipeptides

In HCT-116 cells, a significant increase in gene expression was observed for *CLDN3*, *CLDN7*, and *TJP*-2 (*p* < 0.05) at 2 h post-treatment (Fig. 1). In fact, we also found that *TJP-2* expression increased significantly with P1 (WG, log2FC = 1.85, *p* = 0.035), P2 (VQ, log2FC = 2.4, *p* = 0.0004), and P9 (GV, log2FC = 2.25, *p* = 0.0014) treatments. However, *TJP-1* expression was rather induced at 6 h after P4 (VQ, log2FC = 1.72, *p* < 0.0001), P7 (TF, log2FC = 1.27, *p* = 0.044), P8 (LA, log2FC = 1.66, *p* < 0.00001), and P9 (GV, log2FC = 1.60, *p* < 0.0001) dipeptide treatments. Moreover, following a 6 h treatment with dipeptides, distinct and overlapping gene expression patterns were observed across the cell lines, i.e. VK2 (vaginal cells), HEC-1A (endometrial cells), and HCT-116 (colon cancer cells) (Fig. 1). At 6 h, *CLDN3* expression was upregulated by P7 (TF, log2FC = 2.03, *p* = 0.049) and P8 (LA, log2FC = 2.49, *p* = 0.003) in all cell lines, while *CLDN4* expression was enhanced only in HCT-116 in response to P3 (log2FC = 1.32, *p* = 0.002) and P10 (log2FC = 1.35, *p* = 0.0007), two isoforms of WG dipeptide as well as in response to P5 (LQ), P6 (AL), P7 (TF), and P9 (GV) (log2FC > 1.5, *p* < 0.0001). Hence, we found cell-line-specific responses represented for *CLDN7*, with induced expression in HCT-116 (log2FC = 3.55, *p* = 0.002) and reduced expression in VK2 (log2FC = 0.68, *p* = 0.42) and HEC-1A (log2FC = 0.44, *p* = 0.03) upon P9 treatment for 6 h. Similarly, at 6 h, P1 repressed *CLDN7* in HEC-1A (log2FC = 0.36, *p* = 0.009) but strongly induced its expression in intestinal cells (log2FC = 4.54, *p* < 0.0001). Additionally, *CLDN1* expression was enhanced by P6, P7, P8, and P9 in HEC-1A and HCT-116 at 6 h (*p* < 0.05), whereas in VK2, these dipeptides upregulated the expression of *CLDN1* at 24 h (*p* < 0.05). Equally, *CLDN14* expression increased in HEC-1A (log2FC = 3.23, *p* = 0.0097) and HCT-116 (log2FC = 1.99, *p* < 0.0001) following P5 exposure at 6 h, while in VK2 cells a similar upregulated effect was observed at 24 h (log2FC = 1.61, *p* = 0.62). Likewise, *TJP2* expression was induced at 2 h in HCT-116 (log2FC = 2.4, *p* = 0.0004) and at 6 h in HEC-1A (log2FC = 1.93, *p* = 0.002) by P2 peptide. Similar upregulated levels of *TJP2* were observed with P6 treatment at 6 h in HCT-116 (log2FC = 2.13, *p* = 0.0041) and at 48 h in HEC-1A (log2FC = 3.01, *p* < 0.0001). As positive controls, Vitamin D and lactic acid treatment were included, and they caused a significant increase in the expression of *CLDN1* (*p* < 0.001), *CLDN3* (log2FC > 10, *p* < 0.0001), *CLDN4* (*p* < 0.0001), *TJP1* (log3FC > 6, *p* < 0.0001), *TJP2* (log2FC = 1.8, *p* < 0.0001), and *OCLN* (*p* < 0.0001) expression in VK2 cells.

**Figure 1:**
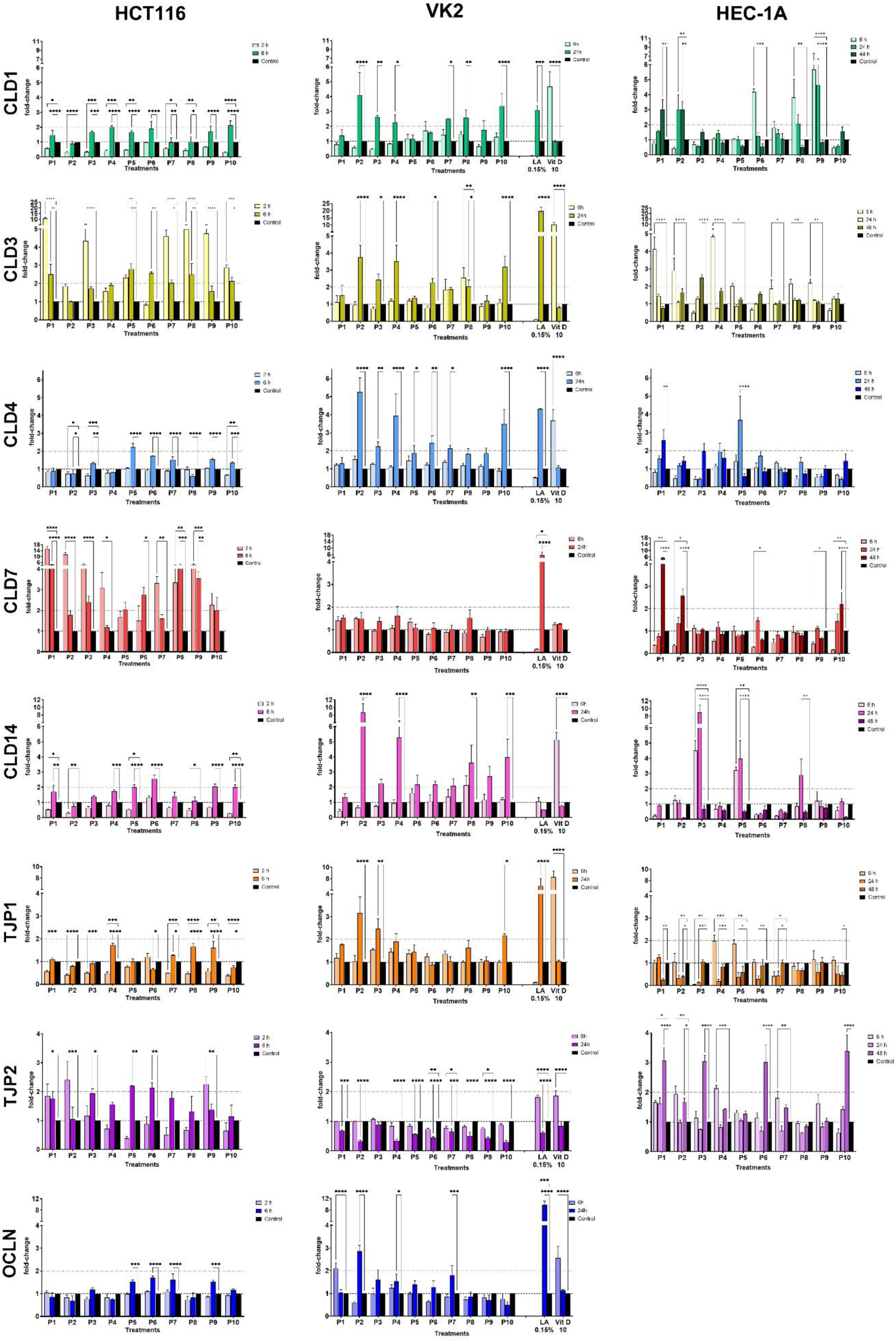
Quantitative PCR (qPCR) analysis of tight junction gene expression across multiple cell lines and treatment conditions. Three human cell lines (VK2, HCT-116, and HEC-1A) were exposed to 10 different dipeptides (Table 1) at various time points. Gene expression levels were normalized to housekeeping genes and are presented as fold change relative to untreated controls. Data represent mean ± SEM. Expression levels are divided horizontally by gene and vertically by cell line. Statistical significance was assessed using one-way ANOVA; *p* < 0.05 was considered significant. * *p* < 0.05; ** *p* < 0.01; *** *p* < 0.001; **** *p* < 0.0001. *P1*: WG-OH; *P2*: VQ-OH; *P3*: WG-NH_2_; *P4*: VQ-NH_2_; *P5*: LQ-NH_2_; *P6*: AL-NH_2_; *P7*: TF-NH_2_; *P8*: LA-NH_2_; *P9*: GV-NH_2_; *P10*: WG-SO_4_.

A comparison of gene expression patterns between VK2 and HEC-1A cells revealed common responses to the tested peptides following 24 h treatments (Fig. 1). Both cell lines exhibited an increase in *CLDN4* gene expression in response to P5 (*p* < 0.05). More specifically, VK2 cells exhibited significant *CLDN4* upregulation after exposure to an array of peptides, such as P2, P4, and P10 (log2FC > 3, *p* < 0,0001) among others (Fig. 1). Additionally, *CLDN3* expression showed a shared increased pattern in HEC-1A cells at 48 h (log2FC = 2.5, *p* < 0.0001) and in VK2 cells at 24 h (log2FC = 2.43, *p* = 0.013) after exposure to P3 (WG), indicating that these peptides can induce similar expression patterns in both cell lines, albeit with variations in duration of exposure. More specifically, VK2 cells exhibited enhanced gene activation after 24 h of dipeptides exposure compared to HCT-116 cells, which portrayed induced expression of tight junction genes at early time of exposure from 2-6 h. In HEC-1A, however, 48 h treatments led to higher expression of TJ genes (Fig. 1). The highest expression level of *TJP2* in HEC-1A was observed at 48 h with P1, P3, and P10 treatments (log2FC > 3, *p* < 0.0001), which are three distinct isoforms of the WG dipeptide. Overall, these data indicate that dipeptides selectively modulate TJ protein expression in a cell-type-dependent manner, with some peptides eliciting consistent effects across multiple epithelial cell types, while others exhibit tissue-specific regulatory roles.

### Dipeptide treatment alters tight junction protein expression and localization

To validate the gene expression findings in VK2 (vaginal epithelia) and HCT-116 (colon cancer) cell lines, we also analyzed the impact on TJ protein expression of different dipeptides through confocal microscopy (summarized in Table 4). Immunofluorescence microscopy followed by mean fluorescence intensity (MFI) quantification revealed distinct expression profiles for CLDN1, CLDN7, and CLDN14 in VK2 cells (24 h) and HCT-116 cells (2 h and 6 h), under specific exposure conditions.

**Table 4:**
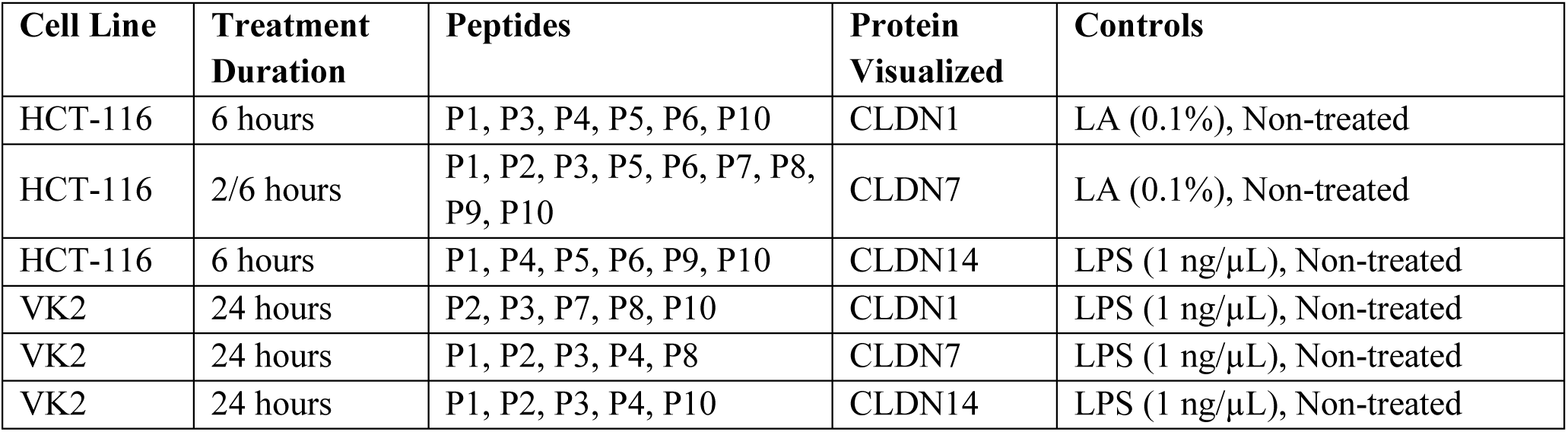
Peptide treatment conditions in this study for immunofluorescence.

In VK2 cells, we observed CLDN1 expression to be significantly increased following treatments with P2 (VQ, *p* < 0.0001) and P3 (WG, *p* = 0.0195) compared with the untreated control at 24 h (Fig. 2 A-B; Fig. S3A). Induced expression of CLDN7 was portrayed post-incubation with P1 (WG, *p* = 0.026) for 24 h (Fig. 2 C-D; Fig. S3B). For CLDN14, no significant changes were detected under any treatment conditions, but a moderate increase was observed after P4 (VQ) and P10 (WG) exposures (Fig. 2 E-F; Fig. S3C). Both CLDN1 (Fig. S3A) and CLDN14 (Fig. S3C) exhibited a cytosolic distribution in VK2. Enhanced expression of CLDN1 was associated with increased protein aggregation in the membranes, as observed following treatments with P2 (*p* < 0.0001), P3 (*p* = 0.0052), P7 (TF), or P8 (LA) (Fig. S3A). In the case of CLDN14 certain treatments such as P10 or LPS caused the formation of a more compact aggregate structure, which were clearly visualized through microscopy (Fig. S3C).

**Figure 2:**
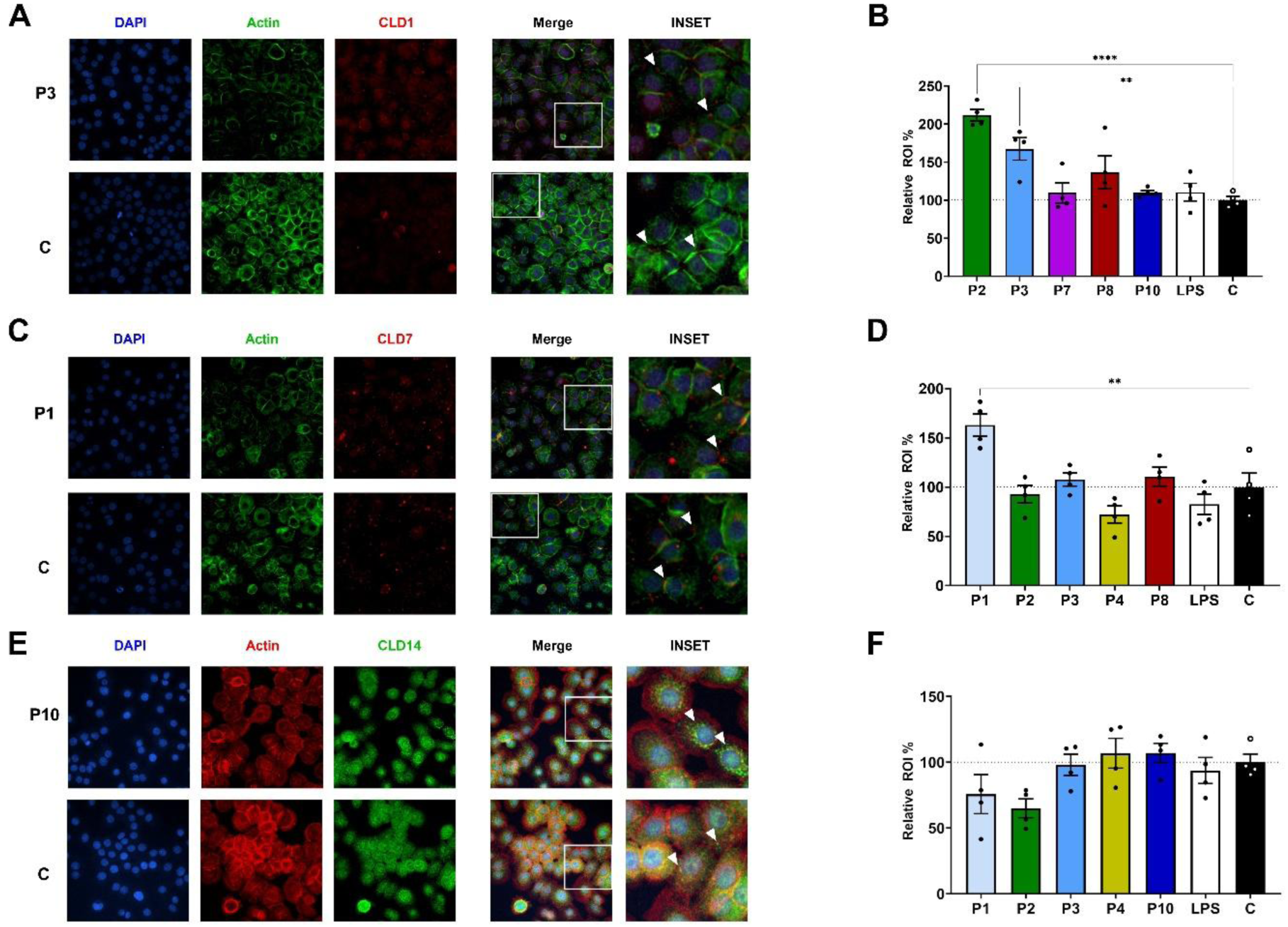
Effect of 24 h dipeptides treatment on claudins’ localization and expression in VK2 vaginal epithelial cells. **A, C, E)** Representative high-resolution confocal micrographs (60X magnification) of VK2 cells showing staining for nuclei (DAPI, blue), F-actin (phalloidin, green or red), and **A**) CLDN1 (red) under P3 (WG) treatment and in untreated control, **C)** CLDN7 (red) under P1 (WG) treatment and in untreated control, or **E)** CLDN14 (green) under P10 (WG) treatment and in untreated control. For each condition, individual fluorescence channels and the merged image are presented. Insets display magnified regions of interest highlighting claudin distribution at the cell–cell junctions, with white arrows indicating enhanced claudin localization. **B, D, F)** Quantification of **B**) CLDN1, **D**) CLDN7, or **F)** CLD14 mean fluorescence intensity (MFI), normalized to DAPI signal to account for cell number. Data are expressed as percentage relative to untreated control (set as 100%). Bars represent the mean ± SEM from 4 independent experiments micrographs. Statistical analysis was performed using one-way ANOVA; *p* < 0.05 was considered significant. ** *p* < 0.01; **** *p* < 0.0001. *P1*: WG-OH; *P2*: VQ-OH; *P3*: WG-NH_2_; *P4*: VQ-NH_2_; *P7*: TF-NH_2_; *P8*: LA-NH_2_; *P10*: WG-SO_4_.

In HCT116, we observed enhanced protein levels of CLDN1 due to P5 (LQ) treatment (*p* = 0.0016) for 6 h (Fig. 3 A-B). Moreover, other dipeptides, such as P1, P3, P4 (VQ), P6 (AL), showed higher median values for CLDN1 expression, although they were non-significant compared with control (Fig. 3B; Fig. S4A). We found that *CLDN7* expression was induced at 2 h, as observed from the gene expression levels *in vitro* (Fig. 1). Although, similar results were also obtained for protein expression, where P3 (*p* < 0.0001) and P9 (GV, *p* = 0.0266) showed significant augmented levels of CLDN7 at 2 h treatment (Fig. 3 C-D; Fig. S4B).

**Figure 3:**
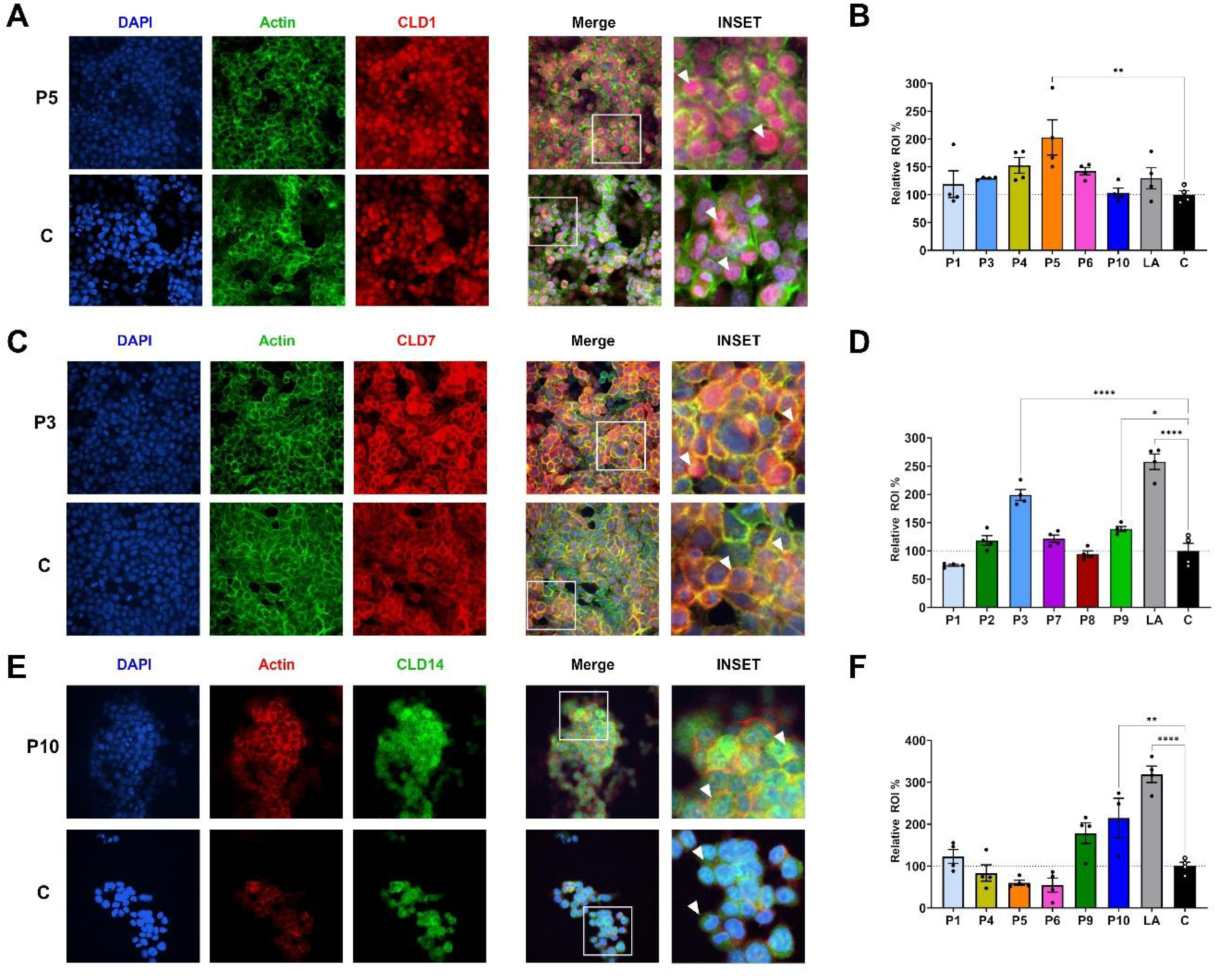
Effect of dipeptides treatment on claudin localization and expression in HCT-116 intestinal epithelial cells. **A, C, E)** Representative high-resolution confocal micrographs (60X magnification) of HCT-116 cells showing staining for nuclei (DAPI, blue), F-actin (phalloidin, green or red), and **A)** CLDN1 (red) under 6h P5 (LQ) treatment and untreated control, **C)** CLDN7 (red) under 2 h P3 (WG) treatment and untreated control, or **E)** CLDN14 (green) under 6 h P10 (WG) treatment and untreated control. For each condition, individual fluorescence channels and the merged image are presented. Insets display magnified regions of interest highlighting claudin distribution at the cell–cell junctions, with white arrows indicating enhanced claudin localization. **B, D, F)** Quantification of **B**) CLDN1, **D**) CLDN7, or **F)** CLD14 mean fluorescence intensity (MFI), normalized to DAPI signal to account for cell number. Data are expressed as percentage relative to untreated control (set as 100%). Bars represent the mean ± SEM from 4 independent experiments micrographs. Statistical analysis was performed using one-way ANOVA; *p* < 0.05 was considered significant. * *p* < 0.05; ** *p* < 0.01; **** *p* < 0.0001. *P1*: WG-OH; *P2*: VQ-OH; *P3*: WG-NH_2_; *P4*: VQ-NH_2_; *P5*: LQ-NH_2_; *P6*: AL-NH_2_; *P7*: TF-NH_2_; *P8*: LA-NH_2_; *P9*: GV-NH_2_; *P10*: WG-SO_4_.

Furthermore, CLDN7 expression was significantly increased after P8 (LA, *p* = 0.0057) exposure at 6 h compared with non-treated cells (data not shown) corroborating with our gene expression findings (Fig. 1). CLDN14 expression was significantly elevated in response to P10 (*p* = 0.007) at 6 h treatments. Although P1 and P9 showed an increase trend for CLDN14 expression, the analyses were not significant compared with the non-treated controls (Fig. 3 E-F, Fig. S4C). Our analysis revealed CLDN1 localization in HCT-116 in the nucleus, whereas CLDN7 localizes in the membrane as observed from Fig. 3 and Fig. S4B. Lactic acid exposure led to significantly increased proteins levels of CLDN7 (*p* < 0.0001) and CLDN14 (*p* < 0.0001) in HCT-116 (Fig. 3C-F). However, under its exposure, we detected induced CLDN1 expression in the membranes and not in the nucleus in HCT-116 (Fig. S4A). In VK2, CLDN7 showed localized expression in the membranes (Fig. S3B), though its presence was increased after some treatments with certain DPs, such as P1.

Additionally, to investigate potential synergistic or cumulative effects, we also tested combinations of dipeptides in our experiments that individually elicited the strongest induction of CLDN1 (Fig. S5A-B) and CLDN14 (Fig. S5C-D) expression in the intestinal cell line, HCT-116. These combinations, however, did not produce significant changes in protein expression levels or subcellular localization compared to non-treated controls (Fig. S5).

### Proteomic and KEGG pathway analysis reveal WG-associated immunomodulatory signatures in HIV-susceptible cells and predicted tight junction interaction networks

Since our *in vitro* data established that EC-derived DPs modulate TJ gene and protein expression in mucosal epithelial models, we subsequently conducted an in-depth analysis of our previously conducted proteomics data (Ceña-Diez et al., 2023) to compare the differential expression of proteins modulated by EC-specific metabolite, WG in HeLa CD4⁺CCR5⁺ cells. Proteomic profiling of WG-treated versus untreated cells identified several proteins associated with inflammatory signaling, interferon response, and cellular stress pathways, hinting at a potential immunomodulatory role of the metabolite WG (Fig. 4, Table S1). Notably, at least five interferon-associated proteins, including two interferon-stimulated genes (ISGs), specifically interferon induced protein 44 and interferon gamma inducible protein 16 (log2FC > 1.9, *p* < 0.0001), were significantly downregulated in WG-treated cells compared to untreated controls (Fig. 4B). Similarly, proteins associated with NF-кB signaling {(i.e., RELB Proto-Oncogene (RELB, *p* = 0.001), TNF-α-induced protein 1 (TNFAIP1, *p* = 0.048), and TNF receptor superfamily member 1A (TNFRSF1A, *p* < 0.0001)}, caspase 1 (CASP1*, p* < 0.0001), IL-32 (*p* < 0.0001), and sequestosome-1 (SQSTM1, *p* < 0.0001) showed significantly reduced abundance in the WG-treated group relative to the control. Reduced expression of these pro-inflammatory proteins might indicate a potential anti-inflammatory effect of the treatment (Franchi et al., 2009; Shin et al., 2013; Hayden and Ghosh, 2014). On the contrary, our study showed upregulation of transforming growth factor β receptor 3 (TGFBR3, *p* = 0.047) and SMAD family member 5 (SMAD5, *p* = 0.045) in the treated group (Fig. 4B). Both these proteins have been shown to play a role in immune tolerance and tissue repair (Bilandzic and Stenvers, 2011; Lin et al., 2024). Furthermore, WG treatment led to the significant upregulation of nuclear factor I B (NFIB, *p* = 0.042), Rho Guanine Nucleotide Exchange Factor 25 (ARHGEF25, *p* = 0.039), RAS-related protein Rab-27A (RAB27A, *p* = 0.002), and 26S proteasome non-ATPase regulatory subunit 1 (PSMD1, *p* = 0.041) (Fig. 4B), all of which play vital roles in cellular differentiation, cytoskeletal regulation, and vesicle transport. (Steele-Perkins et al., 2005; Hsu et al., 2011; Biro et al., 2014; Schweitzer et al., 2016; Xu et al., 2016). Interestingly, our data showed that the WG-treated group also exhibited an increase of CD81 (*p* = 0.002), F11 receptor (*p* = 0.26) and the IL-7 (*p* < 0.0001) compared with the control group. Additional proteins associated with pro-inflammatory signaling were significantly downregulated in treated group, including TNF receptor superfamily member 10A (TNFRSF10A, *p* = 0.006) and stimulator of interferon response cGAMP interactor 1 (STING1, *p* = 0.048), C-X-C chemokine receptor type 4 (CXCR4, *p* = 0.059), mitogen-activated protein kinase-activated protein kinase 2 (MK2; *p* = 0.060), and yes-associated protein 1 (YAP1, *p* = 0.055).

**Figure 4:**
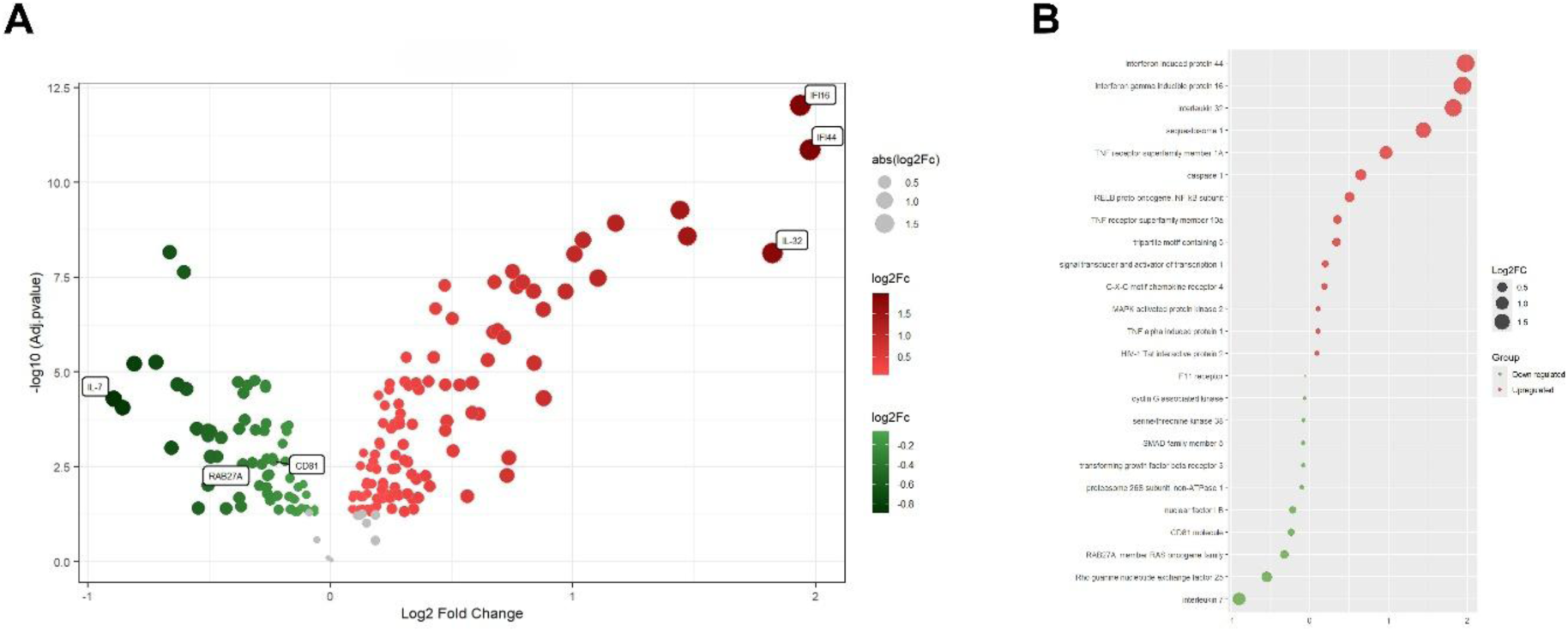
Differential expression analysis of proteins between control and WG-treated HeLa CD4⁺CCR5⁺ cells. **A**) Volcano plot showing the log_2_ fold change (FC) and statistically significant proteins differentially expressed between the control and WG-treated groups. Red and green dots represent proteins upregulated and downregulated, respectively, in the control group compared to the WG-treated group. Three of the most significantly altered proteins in each group are labeled in the figure. **B**) Bubble plot showing a selection of 14 upregulated (red) and 11 downregulated (green) proteins in the untreated control group. Dot size corresponds to the magnitude of log₂ fold change and color indicates direction of change. *p* < 0.05 was considered significant. * *p* < 0.05; ** *p* < 0.01; **** *p* < 0.0001

In line with the individual proteins analysis, KEGG analysis identified significant enrichment of immune and inflammatory signaling pathways involving the differentially expressed proteins (Table S1). NF-κB signaling pathway (hsa04064; *p* = 0.0002), MAPK signaling pathway (hsa04010; *p* = 0.0013), and TNF signaling pathway (hsa04668; *p* = 0.015) were predicted to be downregulated in the treated group. On the contrary, mTOR signaling pathway (hsa04150; *p* = 0.042) was observed to be upregulated in the treated group. RELB, common to NF-κB and MAPK signaling pathways, showed reduced expression in the treated group, as mentioned previously (Fig. 4). NOD-like receptor pathway (hsa04621; *p* = 0.0013) including interferon gamma inducible protein 16 and CASP1, JAK-STAT signaling pathway (hsa04630; *p* = 0.014), and cytokine-cytokine receptor interaction pathway (hsa04060; *p* = 0.0013) were downregulated in dipeptide treated group, whereas PI3K-Akt signaling pathway (hsa04151; *p* = 0.27) showed higher abundance in the treated group.

To further investigate the functional relationships among the TJ genes evaluated by qPCR, predicted protein-protein interaction networks were generated using STRING and merged into a consensus network (Figure S6). STRING network analysis predicted high-confidence interactions among the assessed TJ proteins, supported primarily by co-expression and text-mining evidence. Notably, TJP1 displayed the highest degree of direct interactions in the network, along with TJP2 and OCLN. Interestingly, the merged network also included key proteins such as CD81 and F11 receptor, as well as additional scaffolding proteins associated with TJ complex, like cingulin (CGN), PALS1-associated tight junction protein (PATJ, also known as INADL), and MARVEL domain containing 2 (MARVELD2). CD81 was predicted to interact with both CLDN7 and CLDN1, both of which also showed co-expression with CLDN3. Furthermore, F11 receptor (JAM-A) displayed predicted interactions with CLDN1, CLDN4, CLDN7, TJP2, and OCLN.

## DISCUSSION

Chronic inflammation, immune dysregulation, and epithelial barrier dysfunction are key contributors to many diseases, such as autoimmune disorders, chronic infections, and tissue degeneration (Biro et al., 2014). Chronic mucosal inflammation during HIV-1 infection can, in fact, lead to microbial dysbiosis and translocation, which is associated with systemic inflammation (Tincati et al., 2016). Overall, all these processes can be modulated by microbiome-derived metabolites (Ganieva et al., 2022; Moonwiriyakit et al., 2023). Understanding how these metabolites modulate these processes may have a therapeutic interest. Therefore, applying this rationale to Elite Controllers (ECs) has led to the identification of EC-specific metabolites (i.e. dipeptides WG, VQ, and others) through proteomics (Sperk et al., 2021). Hence, our experimental study investigated the impact of 10 different dipeptides (DPs), metabolites specific to ECs (Sperk et al., 2021), on the expression of tight junction (TJ) proteins in epithelial cells *in vitro*. Our results, interestingly, established a baseline expression profile of TJ genes in different epithelial cell lines for future studies, a previously unexplored subject. Baseline transcriptional profiles differed between VK2 and HEC-1A cells (Fig. S2), including for the absence of detectable *OCLN* expression in the latter. Our data also revealed a tissue-specific response to DP treatment, with distinct TJ proteins expression patterns observed in vaginal, endometrial, and intestinal epithelial cell lines. DP treatment, in fact, varied significantly among the cell types, with the strongest inductions observed in the earliest time period in the intestinal cells, HCT-116 (2 – 6 h), in the intermediate time period in the vaginal VK2 cells (24 h), and in the latest time period in endometrial HEC-1A cells (48 h) (Fig. 1). Remarkably, our findings provide novel insights into the effects of these EC-specific DP treatments enhancing the expression of different TJ proteins in both female reproductive tract and intestinal epithelial models. These *in vitro* results show the potential of these DPs to modulate the epithelial barrier function by regulating claudins and other TJ genes. In fact, our results showed different WG isoforms (which corresponds to P1, P3, or P10 in our experiments, Table 1) enhancing the expression of selective claudins and other TJ genes: *CLDN1, CLDN3, CLDN4, CLDN7, CLDN14*, and *TJP2* in HCT-116; *CLDN1, CLDN3, CLDN4, CLDN14, TJP1*, and *OCLN* in VK2; and *CLDN1, CLDN3, CLDN4, CLDN7, CLDN14,* and *TJP2* in HEC-1A.

Our findings also highlight other key tight junction proteins beyond claudins, including occludin (OCLN), which regulates barrier stability and permeability through PI3K/AKT (phosphatidylinositol 3-kinase and protein kinase B) and MAPK/ERK signaling pathways (Bhat et al., 2019; Marsch et al., 2024), and zonula occludens (ZO/TJP), which links claudins and occludin to the actin cytoskeleton while contributing to cellular signaling and stress responses (Bhat et al., 2019; Citi et al., 2024; Marsch et al., 2024). In HCT-116, P5 (LQ), P6 (AL), and P7 (TF) had significant impact on most of the TJ genes, including *OCLN.* Moreover, at least one of the WG isoforms (P1, P3, P10) enhanced the expression of all the TJ genes except *TJP1* and *OCLN*. However, in VK2 cell line, P2 (VQ) and P10 (WG) showed the most significant effects. In VK2 cells, the highest induction of gene expression was observed for *CLDN3* in all the three isoforms of WG. In contrast, *CLDN7* showed the strongest response to WG in HCT-116. Additionally, in HEC-1A, *CLDN14* exhibited the highest expression in response to the same DP. At the functional level, different claudins regulated by our discovered metabolites play key roles in different signaling pathways. CLDN3 negatively regulates the PI3K/AKT pathway, thereby having a tumor-suppressive function and inhibition of lymphatic endothelial cell migration (Castro Dias et al., 2019; Lei et al., 2022). On the contrary, CLDN7 is regulated through Wnt/β-catenin signaling, and its dysregulation simultaneously disrupts epithelial integrity, impacting inflammatory cascades (Bianchi et al., 2019; Wang et al., 2024). CLD7 also interacts with CLDN1 and CLDN3 to stabilize cellular junctions and suppress tumorigenic pathways like MAPK/ERK (Lu et al., 2011). From microscopy analyses, we observed WG peptides to regulate the induced expression of CLDN7 in both HCT-116 and VK2 cells, as well as upregulate the protein levels of CLDN1 in VK2 cells (Fig. 2), and CLDN14 in HCT-116 (Fig. 3). These dipeptides also induced claudin gene expression in HEC-1A, specifically P1 and P10 for *CLDN7*, P1 for *CLDN1*, and P3 for *CLDN14.* Earlier studies showed CLDN1 to promote epithelial proliferation and inflammation through ERK1/2-dependent Notch signaling (Pope et al., 2014; Tao et al., 2023), while CLDN14, showing the strongest induction in HEC-1A cells in our study, regulates ion transport via PI3K/AKT, potentially activating the Wnt/β-catenin signaling pathway (Bhat et al., 2019; Tao et al., 2023). Altogether, these interactions suggest that EC-derived dipeptides might influence epithelial barrier function through possible modulation of key proteins associated with signaling networks.

In parallel, proteomic profiling of the previously established and characterized antiviral dipeptide, WG, in HeLa CD4^+^ CCR5^+^cell model, generated a significantly altered immune expression profile as observed in our study (Fig. 4, Table S1), suggesting that EC-derived metabolites may likely participate in two distinct processes to control HIV-1 infection: either through a direct effect on the mucosal barrier genes or via the attenuation of inflammatory stress signaling. For instance, our proteomics analyses revealed the upregulation of CD81 in the WG-treated group in cells compared to untreated controls. Given CD81’s established interactions with integrins and tight junction proteins (Gordón-Alonso et al., 2012; Tejera et al., 2013), our finding suggests a possible immunomodulatory effect for epithelial integrity Additionally, the increase in F11 receptor (JAM-A) observed in WG-treated cells in our analysis is also interesting, since JAM-A has been described to have a key role in TJ formation and epithelial integrity (Bonilha et al., 2020), providing insights for further investigation in more complex experimental systems. Our STRING-based *in silico* analysis (Fig. S6), in fact, showed CD81 interacting with both CLDN7 and CLDN1, while the F11 receptor (JAM-A) showed potential interaction with CLDN1, CLDN4, CLDN7, TJP2, and OCLN, as previously observed (Gordón-Alonso et al., 2012; Tejera et al., 2013), identifying these dipeptide-regulated proteins as potential participants of established barrier maintenance pathways. STRING interaction network analysis also predicted functional relationships among several of the TJ proteins evaluated, including several claudins (such as CLD1, CLD7, CLDN3), TJPs, scaffolding proteins (such as MARVELD2, CGN), CD81, and F11R. Beyond the core TJ-associated proteins, the network included MARVELD2 (tricellulin), CGN (cingulin), and PATJ (PALS1-associated tight junction protein), which link TJ complexes and epithelial barrier function to immune signaling pathways discussed above. Tricellulin supports the epithelial barrier and activates NF-κB signaling (Ikenouchi et al., 2005; Zhang et al., 2020) and cingulin binds to TJ proteins and regulates *CLDN2* expression as well as activates MAPK signaling while stabilizing endothelial barriers under inflammatory stress (Guillemot et al., 2012; Su et al., 2025). PATJ, on the other hand, interacts with CLDN1, OCLN, and TJP3 to stabilize apical-lateral TJ structure, which can be disrupted by viral proteins, promoting immune cell infiltration (Michel et al., 2005; Chai et al., 2021).

For the proteomic analysis, WG treatment was associated with reduced expression of several NF-κB pathway components (i.e., RELB, TNFRSF1A) relative to untreated controls, suggesting an attenuation of inflammatory and antiviral-stress signaling pathways upon WG exposure (Hayden and Ghosh, 2014; Huang et al., 2025). Moreover, the upregulation of NFIB and ARHGEF25 in the WG-treated group suggests potentially enhanced epithelial cell junction stability and immune cell coordination (Steele-Perkins et al., 2005; Hsu et al., 2011). Additionally, the modulation of antiviral signaling pathways adds importance to the dipeptide-driven molecular impact in human cells described above. Downregulation of STING1 and YAP1 was observed in the WG-treated group from our analysis. Persistent STING1 activation is associated with autoinflammatory disorders driven by excessive type I interferon signaling and resulting tissue damage (Krawczyk et al., 2023), so its attenuation may reflect a shift away from pathological immune hyperactivation. YAP1, a transcriptional co-activator involved in immune signaling and cell proliferation whose overactivation has been linked to cancer progression and tissue fibrosis (Noguchi et al., 2018; Li et al., 2024), was also downregulated by WG in our study, indicating a potential role for it in limiting fibrotic and tumor-promoting inflammatory processes. Intriguingly, the proteomics analysis also revealed CLDN12 downregulation in the dipeptide-treated group, although our *in vitro* gene expression data indicates a differential regulation of other claudins (*CLDN1, CLDN3, CLDN4, CLDN7*, and *CLDN14*). This could be due to different cell-specific claudin expression profiles in our results with differences in their distribution patterns. Unlike other claudins, like CLDN1, CLDN3, or CLDN7, which are more structural and crucial for tight junction integrity, CLDN12 is more functionally associated with calcium permeability, forming pores to facilitate calcium absorption in the intestinal epithelia (Beggs et al., 2021).

Consistent with the individual protein changes, KEGG pathway level analysis reported, under WG-treatment, an overall shift from inflammatory signaling. Innate immune and inflammatory signaling pathways including NF-κB, MAPK, JAK-STAT, and TNF signaling, are well known to be central in cytokine production, pathogen sensing, and inflammatory response initiation (Hayden and Ghosh, 2014; Soni et al., 2019; Giarratana et al., 2024), and their chronic activation is a recognized hallmark of HIV-1 disease progression. KEGG pathway analysis from our data revealed a relative downregulation of these immune pathways in WG-treated cells compared with untreated ones, suggesting attenuation of chronic immune activation (Table S1). Furthermore, pathways critical for pathogen recognition and inflammation (i.e., Toll-like and NOD-like receptor pathways) (Hearps et al., 2017) were similarly attenuated in the WG-treated cells in our study. At the level of individual mediators, reduced TNFAIP1 expression in treated cells is consistent with a decrease in TNF-driven inflammatory responses (Zhang et al., 2014; Guo and Yuan, 2020), while reduced CXCR4 expression suggests diminished capacity for inflammation-driven immune cell migration to affected tissues (Teixidó et al., 2018). The stress-induced inflammatory role of MK2 further implies that dipeptide treatment may be relevant in conditions associated with chronic immune activation, where sustained MAPK signaling contributes to tissue pathology (Soni et al., 2019).

On the other hand, some pathways upregulated in the WG-treated group included mTOR and PI3K/AKT signaling, which are mostly involved in cellular metabolism, survival, and tissue homeostasis (Sonoki et al., 2017; Giarratana et al., 2024). This upregulation suggests an environment with enhanced tissue repair and metabolic reprogramming that follows attenuation of acute inflammatory signaling. The mTOR and PI3K/AKT pathways are highly relevant in HIV-1 pathogenesis, since they have been implicated in the regulation of T-cell activation and differentiation, and their upregulation may contribute to a state of immune quiescence, a defining phenotype of Elite Controllers. Whether EC status directly drives this metabolic and immunological profile, or it represents one of the several converging mechanisms, still remains to be determined.

Collectively, the molecular evidence supported by literature, as well as by our novel *in vitro* and *in silico* findings, suggests that EC-specific dipeptides participate in multiple interconnected mechanisms relevant to mucosal immunity and barrier integrity, independently to its already proven antiviral properties (Ceña-Diez et al., 2023). Their capacity to modulate claudin expression in a tissue-specific manner, attenuate pro-inflammatory signaling, and affect antiviral pathways, makes them interesting therapeutic candidates for conditions characterized by chronic inflammation, mucosal barrier disruption, and immune hyperactivation, such as autoimmune disorders, long-term viral persistence, and post-viral syndromes. Translating these preliminary findings into clinically relevant ones, however, will require further investigation into more complex experimental systems that better recapitulate the *in vivo* mucosal biology (Clevers, 2016; Noel et al., 2017; Ingber, 2018). Additionally, individual cell line cultures, while valuable for controlled mechanistic analysis, represent a limited *in vitro* model. Co-culture systems incorporating immune and stromal cells, as well as three-dimensional organoid models that recapitulate the architecture and cellular diversity of mucosal tissues are among the next steps for our future aims. What remains to be seen is if these molecular shifts ultimately improve barrier integrity and immune regulation *in vivo*, and how the activity of these dipeptides might be refined for specific clinical contexts.

