## Supplemental figures for "Exploring the molecular function of metabolites identified in Elite Controllers and their role in epithelial integrity and immune regulation"

[1]Department of Medicine Huddinge, CIM-CID, Karolinska Institutet, Stockholm, Sweden
[2]Department of Infectious Diseases, Karolinska University Hospital, Stockholm, Sweden
[3]Current affiliation: Area of Microbiology of School of Medicine. University of Valladolid, Valladolid, Spain.

Running title: **Tight Junctions and Mucosal Integrity in Elite Controllers**

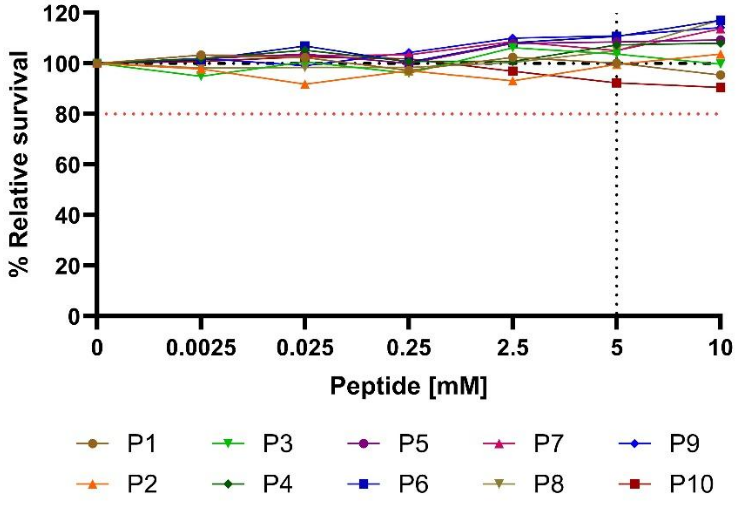


**Figure S1: Cytotoxicity of EC-specific dipeptides.** The cell line HEC-1A was treated with serial dilutions of individual dipeptides up to 10 mM, and after 24 h, the viability was evaluated using CellTiterGlo. Viability of cells that were not exposed to any DP was used for normalization. Black vertical lines indicate 100% and red ones indicate 80% viability.

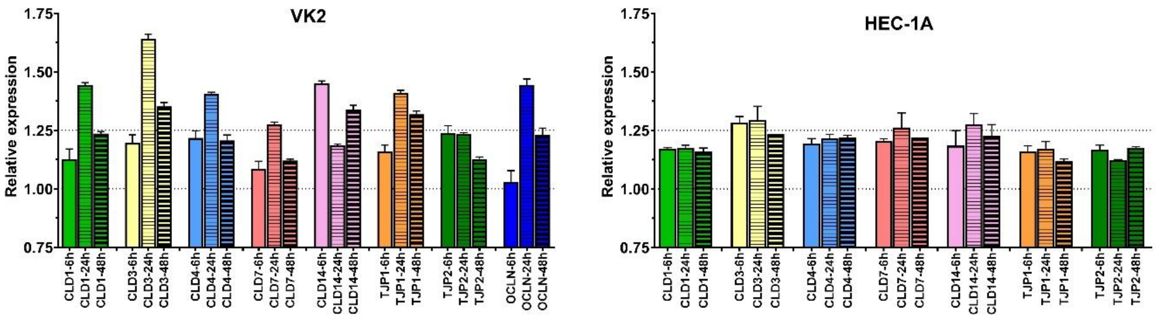


**Figure S2: Baseline expression of tight junction genes in vaginal and endometrial epithelial cell lines.** Quantitative PCR (qPCR) analysis of baseline tight junction gene expression in VK2 and HEC-1A cell lines under untreated conditions. Cells were collected at 6, 24, and 48 hours, and gene expression was assessed for selected tight junction genes based on the housekeeping β-actin gene. Relative expression values are plotted, where lower values indicate higher transcript abundance. Data are presented as mean ± SEM from at least three independent experiments performed in technical triplicates.

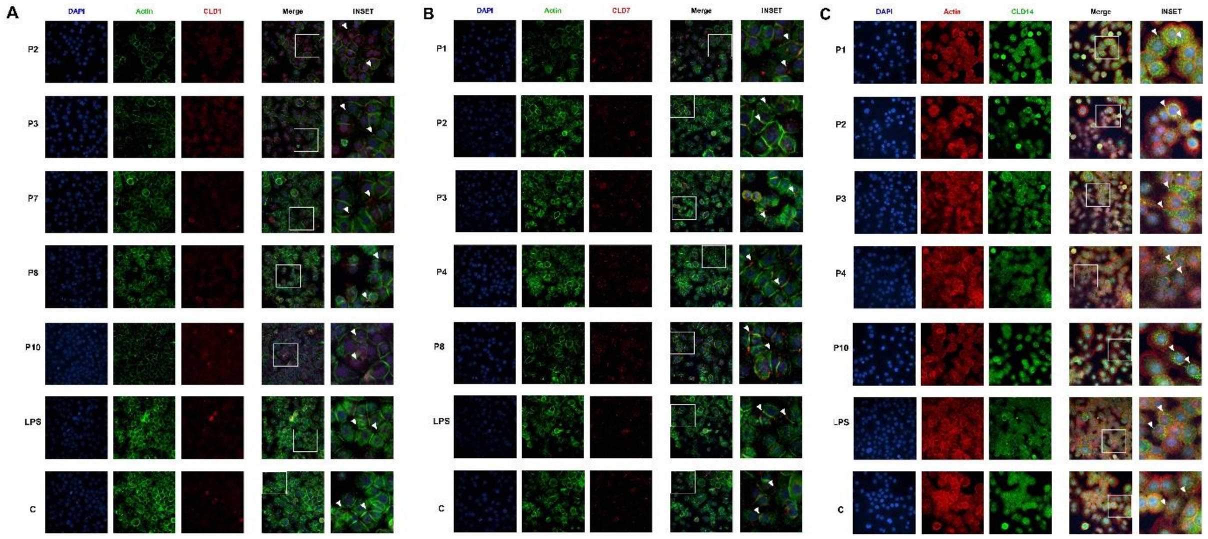


**Figure S3: Effect of 24 h dipeptides treatment on claudins localization and expression in VK2 vaginal epithelial cells.** Representative high-resolution confocal micrographs (60X magnification) of VK2 cells showing staining for nuclei (DAPI, blue), F-actin (phalloidin), and **A)** CLDN1 (red), **B)** CLDN7 (red), or **C)** CLDN14 (green) under different DP treatment and untreated controls. *P1*: WG-OH; *P2*: VQ-OH; *P3*: WG-$NH_2$; *P4*: VQ-$NH_2$; *P7*: TF-$NH_2$; *P8*: LA-$NH_2$; *P10*: WG-$SO_4$.

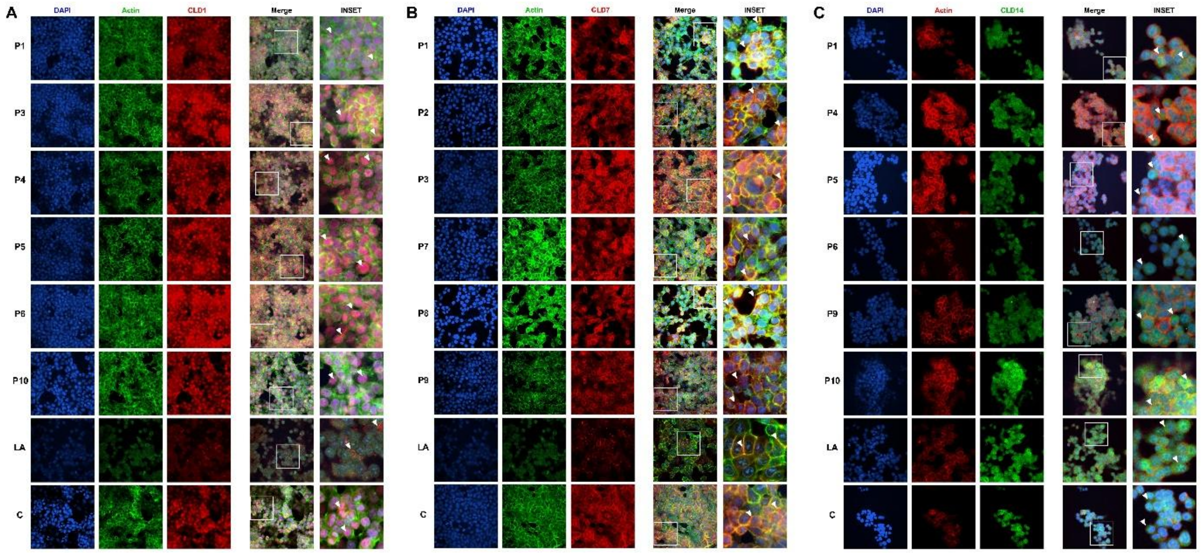


**Figure S4. Effect of dipeptides treatment on claudins localization and expression in HCT-116 intestinal epithelial cells.** Representative high-resolution confocal micrographs (60X magnification) of HCT-116 cells showing staining for nuclei (DAPI, blue), F-actin (phalloidin), and **A)** CLDN1 (red), **B)** CLDN7 (red), or **C)** CLDN14 (green) under **A, C)** 6 h or **B)** 2 h of different DP treatment and untreated controls. *P1*: WG-OH; *P2*: VQ-OH; *P3*: WG-$NH_2$; *P4*: VQ-$NH_2$; *P5*: LQ-$NH_2$; *P6*: AL-$NH_2$; *P7*: TF-$NH_2$; *P8*: LA-$NH_2$; *P9*: GV-$NH_2$; *P10*: WG-$SO_4$.

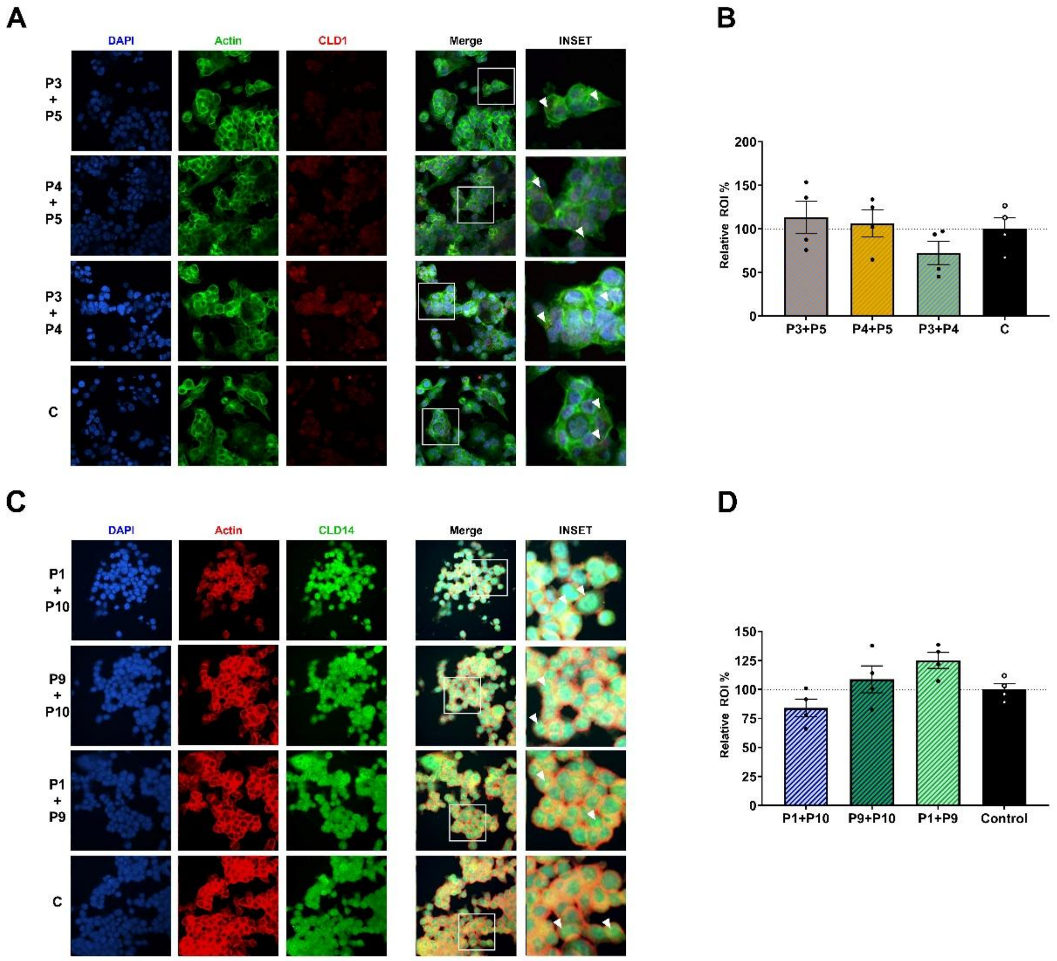


**Figure S5: Effect of 6 h treatment of dipeptide combinations on claudin localization and expression in HCT-116 intestinal epithelial cells. A, C**) Representative high-resolution confocal micrographs (60X magnification) of HCT-116 cells showing staining for nuclei (DAPI, blue), F-actin (phalloidin, red or green), and **A**) CLDN1 (red) under DP combination treatments and untreated control, **C)** CLDN14 (green) under DP combination treatments and untreated control. For each condition, individual fluorescence channels and the merged image are presented. Insets display magnified regions of interest highlighting claudin distribution at the cell-cell junctions, with white arrows indicating enhanced claudin localization. **B, D**) Quantification of **B)** CLDN1 or **D**) CLDN14 mean fluorescence intensity (MFI), normalized to DAPI signal to account for cell number. Data are expressed as percentage relative to the untreated control (set as 100%). Bars represent the mean ± SEM from 4 independent experiments micrographs. Statistical analysis was performed using one-way ANOVA; $p < 0.05$ was considered significant. *P1*: WG-OH; *P3*: WG-$NH_2$; *P4*: VQ-$NH_2$; *P5*: LQ-$NH_2$; *P9*: GV-$NH_2$; *P10*: WG-$SO_4$.

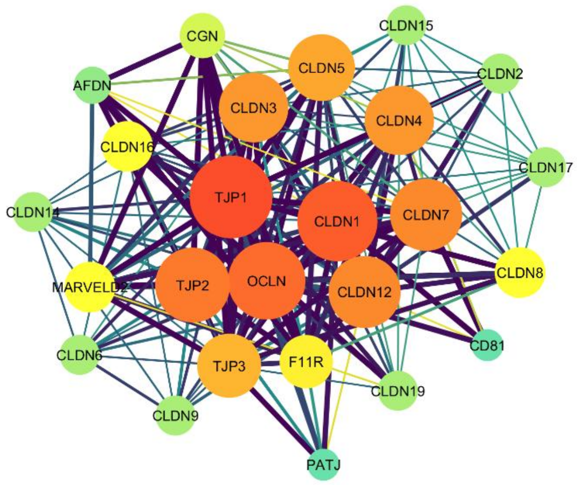


**Figure S6: Merged STRING protein-protein interaction network of TJ and TJ-associated proteins.** Consensus network generated by merging individual STRING interaction networks built for each TJ gene evaluated by qPCR, filtered to show the nodes shared across individual networks. Node color and size are mapped continuously by degree (number of direct interactions per node) from blue, the lowest (2), through yellow, intermediate (14), to highest (26), in red. The edge width is also mapped continuously based on STRING text-timing score (range 0.40-0.99), and the edge color is mapped continuously by STRING combined interaction score (yellow=0.40, blue=0.70, purple=0.99). TJP1 is represented with the highest-degree node (hub) in the network, followed by TJP2, OCLN, and CLDN1, consistent with their central role at the tight junction formation.

**Table S1:** Differentially expressed proteins and pathway enrichment analysis of WG-treated HeLa $CD4^+$ $CCR5^+$ cells compared to control.
